# Nuclear Plasticity Increases Susceptibility to Damage During Confined Migration

**DOI:** 10.1101/2020.01.18.911529

**Authors:** Abhishek Mukherjee, Amlan Barai, Ramesh K Singh, Wenyi Yan, Shamik Sen

**Affiliations:** IITB-Monash Research Academy, Mumbai, India; Dept. of Mechanical Engineering, IIT Bombay, Mumbai, India; Dept. of Mechanical and Aerospace Engineering, Monash University, Clayton, Australia; Dept. of Biosciences & Bioengineering, IIT Bombay, Mumbai, India

**Keywords:** nuclear plasticity, nuclear membrane damage, matrix stiffness, confined migration, finite element analysis (FEA)

## Abstract

Large nuclear deformations during migration through confined spaces have been associated with nuclear membrane rupture and DNA damage. However, the stresses associated with nuclear damage remain unclear. Here, using a quasi-static plane strain finite element model, we map evolution of nuclear shape and stresses during confined migration of a cell through a deformable matrix. Plastic deformation of the nucleus observed for a cell with stiff nucleus transiting through a stiffer matrix lowered nuclear stresses, but also led to kinking of the nuclear membrane. In line with model predictions, transwell migration experiments with fibrosarcoma cells showed that while nuclear softening increased invasiveness, nuclear stiffening led to plastic deformation and higher levels of DNA damage. In addition to highlighting the advantage of nuclear softening during confined migration, our results suggest that plastic deformations of the nucleus during transit through stiff tissues may lead to bending-induced nuclear membrane disruption and subsequent DNA damage.

## Introduction

Cells transit through a myriad of environments, ranging from 2D basement membranes (BM) to 3D collagen networks for morphogenesis, division and proliferation, wound healing and cancer invasion [1]. Cells sense the surrounding mechanical environment to decide on transiting through a pore, a decision that is intrinsically linked to its chances of survival [2]. PDMS devices, widely used for studying confined migration, are significantly stiffer (≈ MPa) than soft tissues (≈ kPa) *in vivo*, thus failing to recapitulate the interplay of nucleus and tissue properties that likely dictates the dynamics of confined migration. The importance of nuclear properties, namely stiffness, in regulating the efficiency of confined migration, is well appreciated. While physical properties of the nucleus are dictated by expression of the intermediate filament protein Lamin (A/C and B) [3–5] and its phosphorylation [6, 7], nuclear deformation is mediated by the actomyosin and the microtubule cytoskeleton which are physically coupled to the nucleus via nesprins [8, 9]. Additionally, localized cytoplasmic stiffening at sites of increased stress from the external environment [10–12], might facilitate nuclear compression thereby aiding in confined migration.

Computational modeling of cell migration has primarily been achieved either by idealizing them as solid continuum spring/spring-dashpot models [13–15] or as liquid droplets bounded by deformable membranes [16, 17]. These assumptions are reasonable in light of a cell being biphasic, exhibiting solid-like behaviour in certain situations and liquid-like character in others. Both of these types of models have their limitations; whereas solid continuum models are unable to replicate similar levels of extreme cellular deformation that occurs in-vivo, liquid droplet models are unable to quantify intracellular stresses. Appropriate visualization of the evolution of stresses within cellular structures such as the actin cytoskeleton and nucleus is critical to complement experimental observations. The current state-of-the-art experimental procedures are unable to predict the stresses that the nucleus undergoes while migrating through 3D confined environments. Traction force microscopy that calculates the stress on the surface of a substrate by relating the deformation of that surface to the stress using Hooke’s law is a critical tool for visualizing mechanical interaction between a cell and its environment, but is limited by its inability to quantify intracellular stresses.

Extreme nuclear deformations during migration through 3 *µ*m transwell pores have been shown to cause plastic deformation [18, 19] as well as nuclear membrane rupture [20–22]. Whether or not nuclear plasticity and nuclear damage are inter-related remains unknown. Also, the extent to which tissue properties influence the plastic deformation of the nucleus has not been probed. For probing nuclear deformation and deformation-induced damage, here we have developed a plane strain finite element model to simulate confined cell migration through a tissue-mimetic environment where mechanical properties of the cell and nucleus have been considered. Studying the collective influence of nuclear and tissue stiffness on the dynamics of pore migration, our results predict the magnitude of cellular force required to squeeze through a constriction and the intracellular stresses sustained by the cell. Our results predict that stiff nuclei passing through stiffer tissues undergo plastic deformations leading to nuclear membrane bending, which may be the cause of nuclear rupture documented experimentally. We validate these predictions using experiments wherein nuclear stiffening led to plastic deformation of the nucleus and higher DNA damage. In addition to predicting a scaling relationship between the timescales and force-scales associated with pore entry, our results establish a direct link between nuclear plasticity and nuclear damage during constricted migration.

## Results

### Nuclear and tissue properties collectively dictate dynamics of confined migration

The nucleus which is the largest and stiffest organelle inside the cell, is physically connected to the cytoskeleton through the LINC complex [23]. Consequently, compression of the cell during confined migration is associated with compression of the nucleus with the extents of cytoplasmic/nuclear deformations dictated by their mechanical properties in relation to that of the surrounding tissues. For studying dynamics of confined migration, a finite element model was developed wherein physical properties of cell membrane, cell cytoplasm, and nucleus were taken into account. Consistent with experiments, the cell membrane, cell cytoplasm and nuclear membrane were modeled as viscoelastic Kelvin-Voigt materials (Fig. S1; refer to Computational Methods section and Supplementary Information for details) [24–26]. A similar viscoelastic description was also used in modeling tissue behavior [27]. Furthermore, consistent with stress-induced permanent deformation of nucleus, an elastoplastic behavior was assumed [28, 29]. Finally, cytoskeletal strain stiffening behavior observed with reconstituted cytoskeletal networks was also accounted for [30–32].

In our model, cell migration through pores in tissues/deformable matrices was assumed to be frictionless and mediated by protrusive forces exerted at the leading edge (Fig. 1a). To first probe the effect of nuclear size on migration efficiency, simulations were performed wherein dynamics of cell entry into a pore of given size (i.e., *ϕ* = {3, 5} *µ*m) was tracked for different sizes of nucleus (i.e., *D*_0_ = 5 and 6 *µ*m) and for varying tissue stiffness (i.e., *E*_*T*_ : (500 − 2000) Pa) (Fig. 1b). In these simulations, nuclear stiffness was kept constant at *E*_*n*_ = 1 kPa. For entry into a pore within the same tissue, i.e., *E*_*T*_ = *E*_1_ = *E*_2_, the time for pore entry (*T*_entry_) as well as the maximum force required for pore entry (*F*_entry_) remained unchanged irrespective of *E*_*T*_ when the nucleus was smaller or equal to the pore size (i.e., *D*_0_*/ϕ* ≤ 1) (Fig. 1c). However, both these quantities increased with increase in *E*_*T*_ for *D*_0_*/ϕ* > 1, highlighting the role of the nucleus in regulating confined migration. When *E*_1_ ≠ *E*_2_, which corresponds to entry at the interface of two distinct tissues, *T*_entry_ and *F*_entry_ were comparable to values corresponding to the higher tissue stiffness (Fig. 1d).

**Figure 1:**
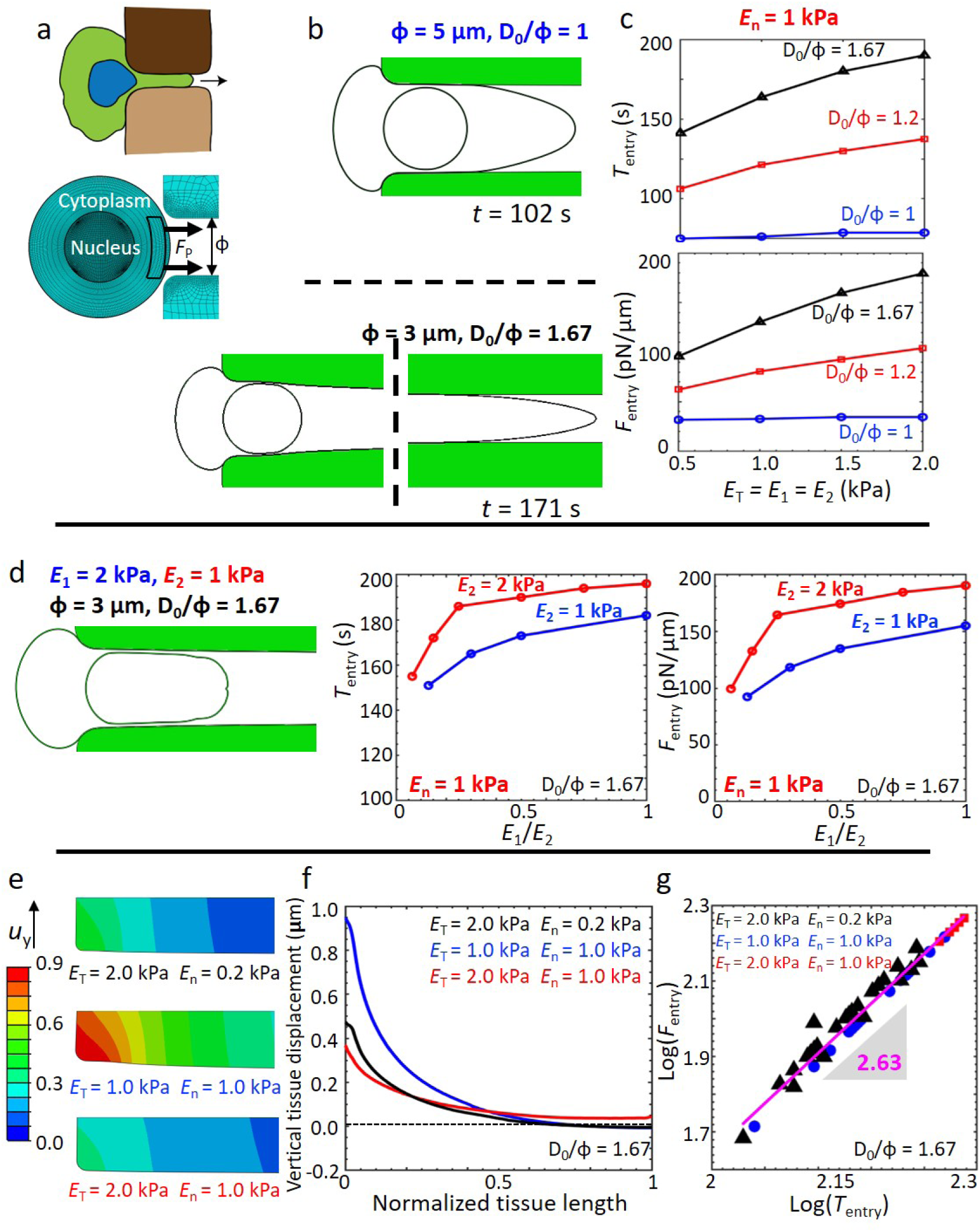
Interplay of nuclear and tissue stiffness on dynamics of pore entry. **a**, Schematic of a cell squeezing through a pore in a given tissue or at the interface of two different tissues. **b**, Cellular deformation just after entry into pore for different values of degree of confinement (*D*_0_*/ϕ*). *E*_*c*_ and *E*_*n*_ were kept constant at 1 Pa and 1 kPa, respectively. **c**, Force (*F*_entry_) and time (*T*_entry_) required for a cell to enter a pore of given size and their dependence on tissue stiffness (*E*_*T*_ = *E*_1_ = *E*_2_) and *D*_0_*/ϕ*. **d**, Nuclear deformation for the case of cell entry through an interface between two dissimilar tissues. Dependence of *F*_entry_ and *T*_entry_ on *E*_1_*/E*_2_ for *D*_0_*/ϕ* = 1.67 and *E*_*n*_ = 1 kPa. **e**, Contour plots of vertical tissue displacement (*u*_*y*_) at the time of nucleus entry into the pore, i.e., when the entire nucleus has completed entering the pore. **f**, Spatial dependence of *u*_*y*_ along the tissue length at the time of pore entry for different values of *E*_*T*_ and *E*_*n*_ and *D*_0_*/ϕ* = 1.67. **g**, Scaling relationship between *F*_entry_ and *T*_entry_ for *D*_0_*/ϕ* = 1.67.

Entry into small pores (*D*_0_*/ϕ* = 1.67) was mediated by widening of the pores as evident from the vertical displacement of the tissues in a *E*_*n*_-dependent manner (Fig. 1e). While displacements far from the pore entry decayed to zero in most cases, for the case corresponding to *E*_*T*_ = 2 kPa, *E*_*n*_ = 1 kPa, vertical displacement of the tissue was non-zero even at distances far from the entry point. The maximum vertical tissue displacement exhibited a non-monotonic dependence on *E*_*n*_*/E*_*T*_ with lowest displacement corresponding to *E*_*T*_ = 2 kPa, *E*_*n*_ = 1 kPa where non-zero displacements were observed far from the entry point (Fig. 1f). Plotting of *F*_entry_ versus *T*_entry_ corresponding to *D*_0_*/ϕ* = 1.67 for different combinations of *E*_*T*_ and *E*_*n*_ revealed a nearly cubic scaling relationship with a factor of 2.63 (Fig. 1g). Together, these results suggest that pore migration through deformable matrices is collectively dictated by nucleus and tissue properties with entry time-scales and force-scales strongly coupled to each other.

### Degree of confinement and nuclear/tissue properties collectively dictate average cell speed

To probe how nuclear/tissue properties and the extent of confinement influence cell motility, cell velocity (*v*_*x*_) was tracked along the direction of migration, i.e., *x*-direction. *v*_*x*_ remained nearly zero for an extended duration, and shot up drastically towards the end (Fig. 2a). The dependence of the average velocity ⟨*v*_*x*_⟩ on *E*_*n*_*/E*_*T*_ was dictated by *D*_0_*/ϕ* and *E*_*n*_ (Fig. 2b). Interestingly, ⟨*v*_*x*_⟩ was nearly independent of *E*_*n*_*/E*_*T*_ for *D*_0_*/ϕ* = 1 for all values of *E*_*n*_, as well as *D*_0_*/ϕ* = 1.67 and *E*_*n*_ ≥ 1 kPa. However, for *D*_0_*/ϕ* = 1.2, ⟨*v*_*x*_⟩ scaled positively with *E*_*n*_*/E*_*T*_ and negatively with *E*_*n*_.

**Figure 2:**
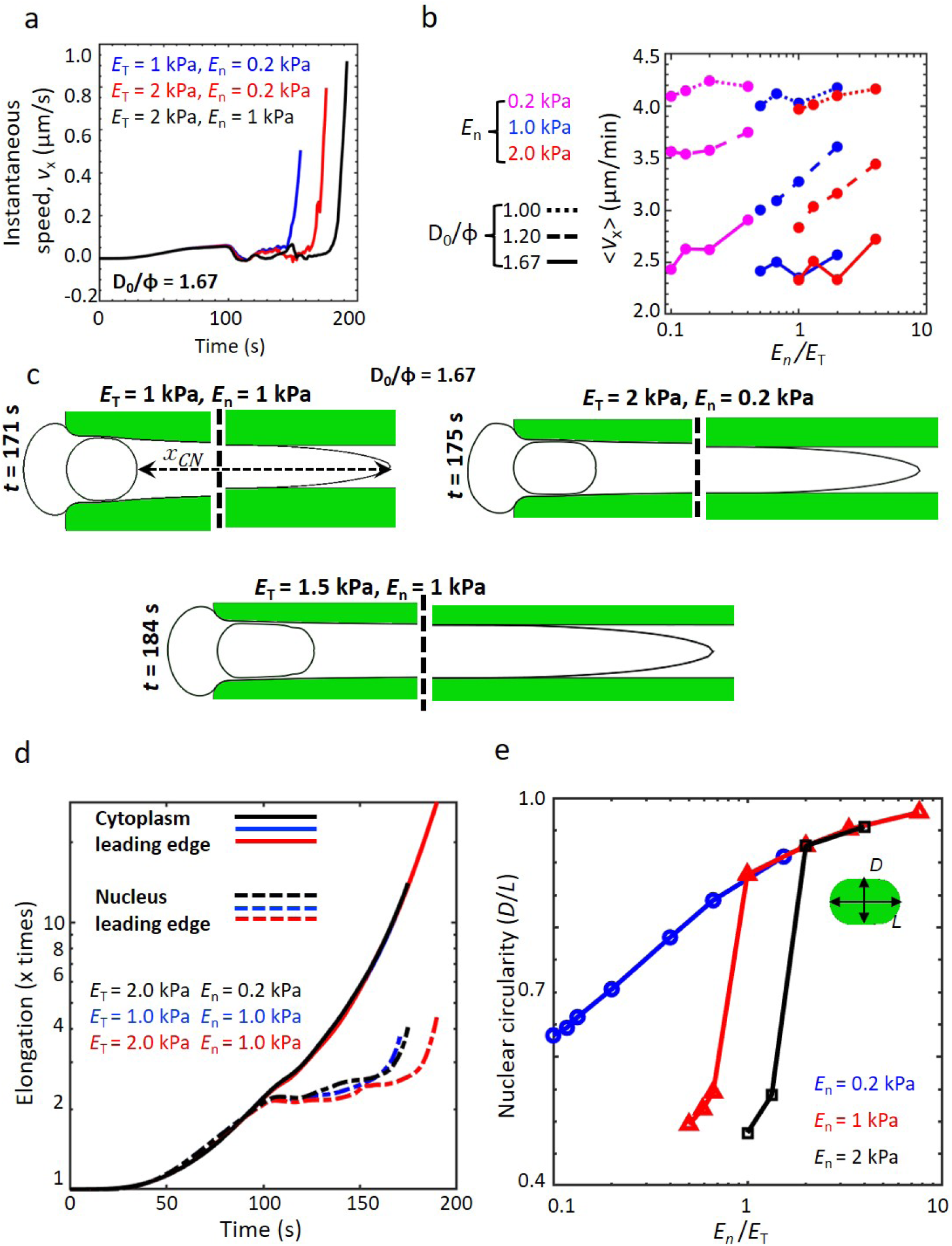
Morphological changes in cell/nucleus during confined migration. **a**, Instantaneous cell velocity (*v*_*x*_) calculated from the start of the simulation (*t* = 0 s) till the instant of pore entry. **b**, The dependence of average cell velocity (⟨*v*_*x*_⟩) on *E*_*n*_*/E*_*T*_ for different values of *D*_0_*/ϕ*. **c**, Shape of the cell and the nucleus at the time of pore entry for different combinations of *E*_*T*_ and *E*_*n*_ and *D*_0_*/ϕ* = 1.67. **d**, Temporal evolution of cytoplasmic and nuclear elongation for *D*_0_*/ϕ* = 1.67. **e**, Dependence of nuclear circularity (*D/L*) on *E*_*n*_*/E*_*T*_ for different values of *E*_*n*_.

Tracking of the normalized distance between the leading edge of the cell and the proximal edge of the nucleus (*x*_*CN*_) revealed upto 30-fold increase in *x*_*CN*_ (Fig. 2c, d). While the period of near zero velocity coincided with duration of extension of the cell cytoplasm with negligible nuclear deformation, the sudden increase in cell velocity (*t* ≈ (160 − 180) sec) corresponded to nuclear entry into the pore. Strikingly, the extent of nuclear deformation, quantified by measuring nuclear circularity (i.e., *D/L*), did not exhibit any dependence on *E*_*n*_ for *E*_*n*_*/E*_*T*_ ≥ 2 (Fig. 2e). However, the individual curves diverged for different values of *E*_*n*_ when *E*_*n*_*/E*_*T*_ < 2. Lowest *D/L*(≈ 0.5) was observed for *E*_*n*_ = {1, 2} kPa and *E*_*n*_*/E*_*T*_ ≈ 1. Together, these results suggest that cell speed is dictated not only by nuclear/tissue properties, but also by the extent of confinement.

### Plastic deformation of the nucleus and kink formation during pore entry

Alteration in nuclear circularity during pore entry is indicative of varying extents of nuclear stresses during and after entry (Fig. 3a, S2). Highest stress in the nucleus was observed for the case of *E*_*n*_ = 0.2 kPa, *E*_*T*_ = 2 kPa, where |*E*_*T*_ − *E*_*n*_| is maximum. Surprisingly, when the nucleus was 5 times stiffer (i.e., *E*_*n*_ = 1 kPa), stresses in the nucleus was lower, and the nucleus was more elongated, raising the possibility of its plastic deformation. Indeed, plastic deformation was observed for cases wherein the nucleus was stiff (i.e., *E*_*n*_ = 1 − 2 kPa) and the tissue stiffer (i.e., *E*_*T*_ > *E*_*n*_) (Fig. 3b). For these cases, dramatic drop in nuclear circularity was observed, i.e., *D/L* < 0.6. Plastic nuclear deformation was associated with reduced nuclear stresses (Fig. 3c), as well as stresses on the the cell membrane (Fig. S3).

**Figure 3:**
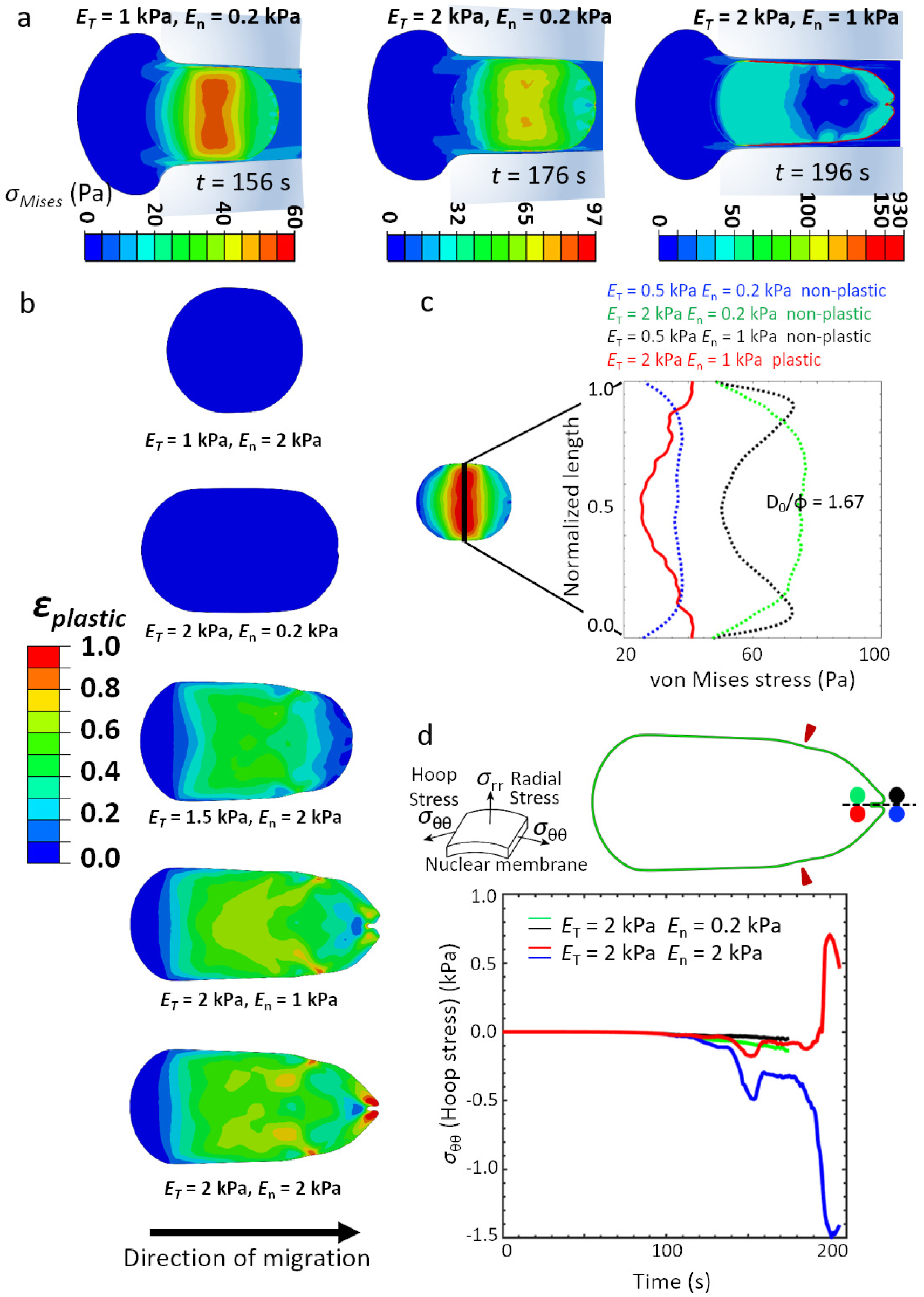
Nuclear plasticity during confined migration. **a**, The spatiotemporal evolution of stress distribution just after entry of the 5*µ*m nucleus into a 3*µ*m pore, i.e., *D*_0_*/ϕ* = 1.67. Contours and colourbars indicate von Mises stresses developed in the cytoplasm and nucleus. **b**, Spatial map of plastic strain accumulated in the nucleus just after pore entry. **c**, Spatial distribution of von Mises stress in the nucleus along the vertical direction just after nuclear entry (*D*_0_*/ϕ* = 1.67). **d**, Temporal evolution of hoop stresses (*σ*_*θθ*_) in the nuclear membrane from the start of simulation to the instant the nucleus completely enters the pore. The two cylindrical components of stresses, namely, radial (*σ*_*rr*_) and hoop (*σ*_*θθ*_) stress in the nuclear membrane are depicted along with the region of nuclear membrane from which the curves are extracted (*D*_0_*/ϕ* = 1.67). Green-Black and Red-Blue curves correspond to two different combinations of *E*_*T*_ and *E*_*n*_ as shown. Green and Red dots in the representative snap-shot of the nuclear membrane correspond to kinked mesh elements on the nuclear membrane at the interface of nucleus and nuclear membrane for *E*_*n*_ = 0.2 kPa and *E*_*n*_ = 2 kPa respectively. Similarly, Black and Blue dots correspond to kinked mesh elements on the nuclear membrane at the interface of cytoplasm and nuclear membrane.

Interestingly, profiles of plastically deformed nuclei revealed the presence of kinks at the front edge with maximum kink formation observed for the case of *E*_*n*_ = 2 kPa, *E*_*T*_ = 2 kPa (Fig. 3b, d). Interfacial migration (i.e., *E*_1_ ≠ *E*_2_) through stiff matrices was also found to be facilitated by plastic deformation (Fig. S4). A plot of temporal evolution of hoop stress (*σ*_*θθ*_) during pore entry revealed varying stress profiles across the front end of the nuclear membrane marked by the green-black and red-blue dots (Fig. 3d). For a stiff nucleus, necking was observed at the lateral edges when it is squeezed to enter the pore (Fig. S5). This was also observed to be the location of initiation of plastic deformation. Necking temporally precedes kink formation at the front edge of the nucleus. For soft nucleus, i.e., *E*_*n*_ = 0.2 kPa, the front end of the nuclear membrane (i.e., green-black dots) was under compressive stresses (negative hoop stress) only. In contrast, for stiff nucleus, i.e., *E*_*n*_ = 1 kPa, while the outer edge of the front end of the nuclear membrane (i.e., blue dot) underwent drastic increase in compressive stresses, the inner edge of the nuclear membrane (i.e., red dot) underwent a sudden switch from compressive to tensile stresses. A switch in stresses in the membrane creates a condition of extreme bending deformations which might be indicative of localized structural disintegration. In addition to highlighting the prominent role of plastic deformation of the nucleus in enabling entry into small pores, our results suggest that buildup of stresses during entry may lead to nuclear membrane damage.

### Nuclear plasticity and DNA damage: insights from experiments

To test our simulation predictions with experiments, confined migration experiments were performed using HT-1080 fibrosarcoma cells which are highly invasive and are capable of switching from proteolytic to non-proteolytic migration upon inhibition of protease activity [33]. This switch is enabled by nuclear softening through phosphorylation of Lamin A/C, and can also be induced by treatment with the non-muscle myosin II inhibitor blebbistatin (hereafter Blebb) [6]. To assess the importance of nuclear stiffness and nuclear plasticity during confined migration, experiments were performed in the presence of Blebb and the CDK inhibitor RO-3306 (hereafter RO), which inhibits lamin A/C phosphorylation [34]. Cells treated with DMSO served as controls. At the drug doses used, no obvious differences in cell morphology were observed (Fig. 4a). While nuclear volume was preserved across the three conditions (Figs. 4b, c), AFM probing of nuclear stiffness with a stiff tip right at the center of the cell (above the nucleus), and fitting of ∼ 2 *µ*m of force curves revealed reduction in nuclear stiffness of Blebb-treated cells compared to controls (Fig. 4d, e). In comparison, RO-treated nuclei were significantly stiffer.

**Figure 4:**
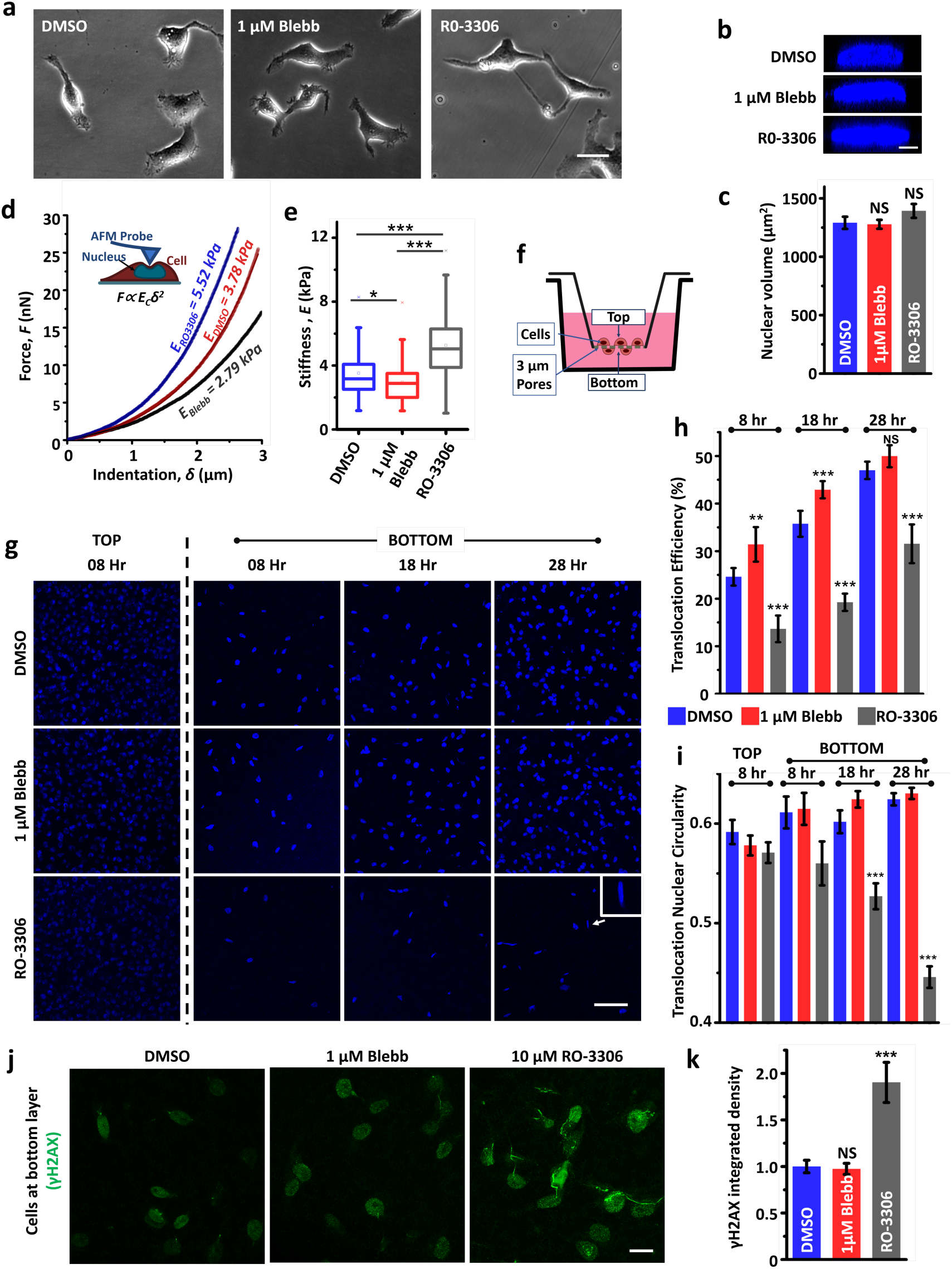
Nuclear plasticity increases susceptibility to DNA damage. **a**, Phase contrast images of HT-1080 fibrosarcoma cells treated with vehicle (DMSO), 1 *µ*M blebbistatin (Blebb) or 10 *µ*M RO-3306 (RO). **b**, Representative XZ plane images of DAPI stained nuclei of DMSO, Blebb and RO-treated cells. Scale bar = 30 *µ*m. **c**, Quantitative analysis of nuclear volume (*n* = 20 − 50 nuclei per condition across 2 independent experiments). Error bars represent ±SEM. Statistical significance was determined by one-way ANOVA/Fisher Test; NS: *p* > 0.05. **d**, Probing nuclear stiffness of cells with a stiff pyramidal probe. Nuclear stiffness values were estimated by fitting 2 ≥ *µ*m of indentation data using Hertz model. **e**, Quantification of nuclear stiffness of DMSO-treated, Bleb-treated and RO-treated cells (*n* = 40−60 nuclei per condition across 2 independent experiments). Error bars represent ±SEM. Statistical significance was determined by one-way ANOVA/Fisher Test; ^*^ *p* < 0.05, ^***^ *p* < 0.001. **f**, Schematic of transwell migration assay through 3 *µ*m pores; Cells were seeded in the upper chamber containing plain DMEM supplemented with DMSO or drugs. Lower chamber was labelled with DMEM containing 20% serum for creating a chemokine gradient. **g**, Representative DAPI stained images of nuclei in upper chamber (referred as TOP) and lower chamber (referred as BOTTOM) at 8, 18 and 28 hrs after cell seeding. **h** (top), Quantification of translocation efficiency of DMSO/Bleb/RO-treated cells at 3 different time-points (*n* 900 nuclei per condition were counted in the upper chamber; experiment was repeated thrice). Error bars represent ±SEM. Statistical significance was determined by one-way ANOVA/Fisher Test; *** *p* < 0.001, ** *p* < 0.01, NS: *p* > 0.05. **i** (bottom), Quantification of nuclear circularity of DMSO/Bleb/RO-treated cells at the top (8 hr time-point) and at the bottom surface of the pores at 3 different time-points (*n* > 80 nuclei per condition; experiment was repeated twice). Error bars represent ±SEM. Statistical significance was determined by Mann-Whitney test; *** *p* < 0.001. **j**, Representative *γ*H2Ax-stained nuclei of DMSO/Bleb/RO-treated cells that have transited to the bottom surface of the pores 28 hrs after cell seeding. **k**, Analysis of integrated intensity of *γ*H2Ax staining in DMSO/Bleb/RO-treated cells (*n* = 40 − 120 nuclei per condition; experiment was repeated twice). Error bars represent ±SEM. Statistical significance was determined by one-way ANOVA/FisherTest; *** *p* < 0.001, NS: *p* > 0.05.

To assess the implications of these alterations in nuclear stiffness vis-a-vis confined migration, transwell migration through 3 *µ*m pores were performed wherein cells were plated on the top of the transwell pores and the fraction of cells reaching the bottom was quantified at three different time-points, i.e., 8, 18 and 28 hours after seeding (Fig. 4f). Cells were stained with DAPI for ease of cell counting as well as for assessing nuclear morphology before and after transit through the pores. Time-snaps of the number of cells that transited through the pores and reached the bottom surface illustrated the clear advantage of nuclear softening during confined migration. While the number of nuclei at the bottom were comparable in DMSO and Blebb-treated cells at all the three time-points, the number of RO-treated nuclei were significantly lesser (Fig. 4g). Quantification of translocation efficiency, i.e., the fraction of cells that transited through the pores, revealed Bleb-treated cells which possessed the softest nuclei to be the most efficient in pore migration, and RO-treated cells to be the least efficient (Fig. 4h).

To next assess the possibility of nuclei undergoing plastic deformation during pore migration, nuclei shape was quantified by measuring nuclear circularity as a function of time. Nuclear circularity of DMSO and Bleb-treated cells remained unchanged across the three time-points, and were comparable with those of cells that remained at the top surface (Fig. 4i). In comparison, nuclear circularity of RO-treated cells dropped significantly as a function of time with a drop of ≈ 25% circularity in cells at 28 hour time-point. The dramatic change in nuclear circularity of RO-treated cells suggests that nuclei of these cells have undergone plastic deformation and retain their deformed shapes. To next probe the link between elastic versus plastic nuclear deformation and nuclear damage, cells were stained with *γ*H2Ax, a marker of DNA damage (Fig. 4j). Quantification of *γ*H2Ax intensity revealed comparable levels in DMSO and Bleb-treated cells, but twice as high in RO-treated cells (Fig. 4k). Together, these results validate our model predictions and suggest that plastic deformation of the nucleus increases susceptibility to DNA damage.

### Scaling relationships

The cellular force required for a nucleus to successfully enter a pore is expected to depend on both nuclear stiffness and tissue stiffness. The dynamic change in nuclear circularity over the period of entry into the pore is then a function of the aforementioned factors. The ratio of initial nuclear size to initial pore size (*D*_0_*/*Φ) is of limited value for analyzing cell migration through deformable matrices because the pore size widens with the passage of a cell nucleus through it. A non-dimensionalized scaling relationship between nuclear circularity (*D/L*) and a combination of tissue and nuclear stiffness shows the slopes followed by cells of varying nuclear stiffness (Fig. S6a). Force required by a cell of given cell/nuclear stiffness for entering a pore is well fit by the following power law with an exponent of 0.5 (equation (1)) (Fig. 5a):

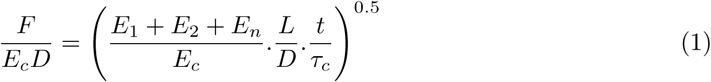

where, *E*_*c*_, *t* and *τ*_*c*_ refer to cytoplasmic stiffness, time of pore entry and viscoelastic time constant of the cytoplasm respectively. The dimensional force itself was found to increase exponentially with *D/L* with the exponents dictated by *E*_*n*_ and the extent of plastic deformation (Fig. S6b). The magnitude of the exponent was highest for the case of stiff nucleus (*E*_*n*_ = 1 kPa) undergoing plastic deformation, lowest for the case of stiff nucleus undergoing non-plastic deformation, and intermediate for the case of soft nucleus (*E*_*n*_ = 0.2 kPa) undergoing non-plastic deformation. These scaling relationships can be utilized for predicting cell generated forces based on experimentally observed parameters such as nuclear circularity and mechanical properties of various cellular and tissue structures.

**Figure 5:**
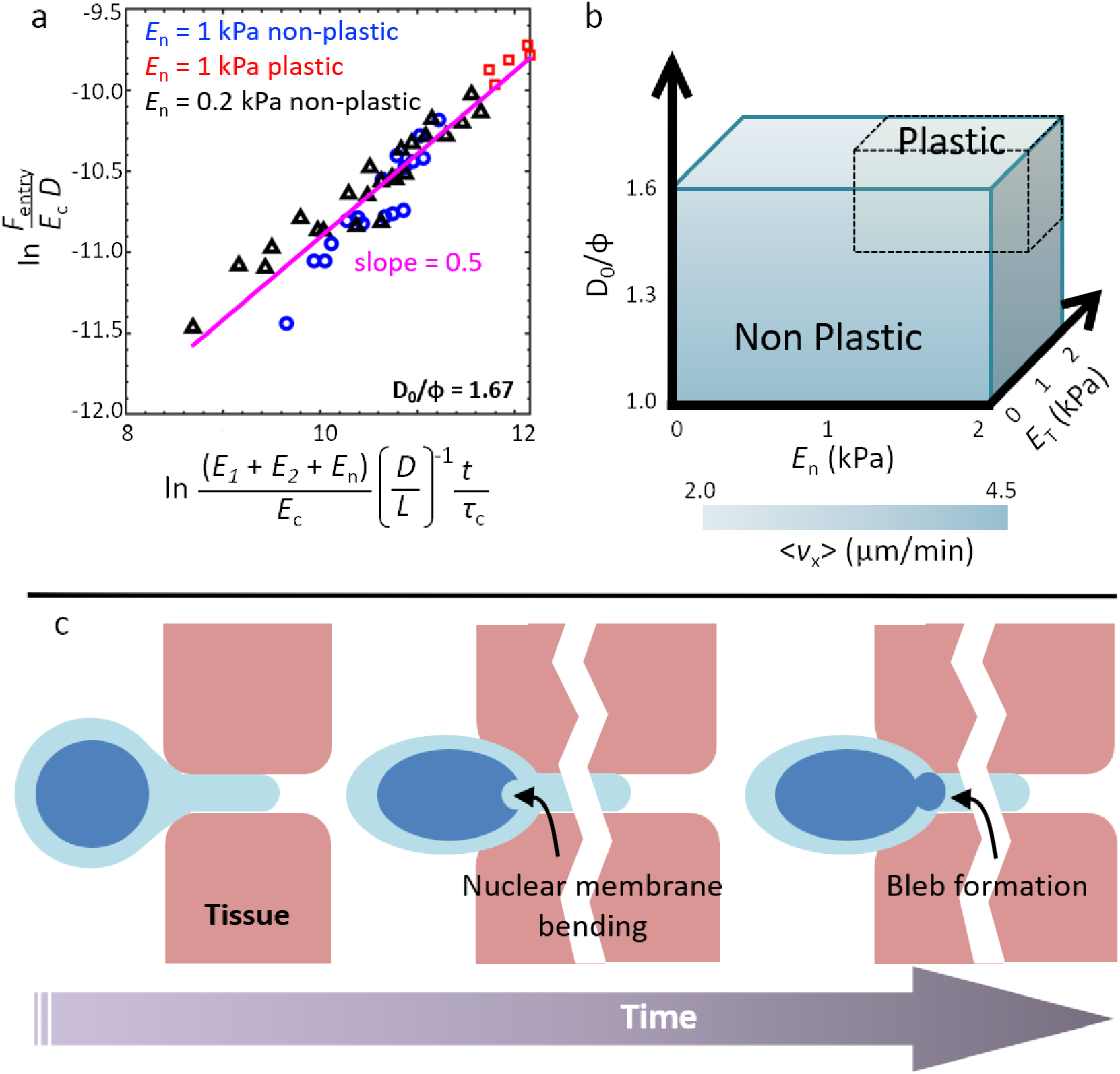
Scaling relationships and proposed model of nuclear damage. **a**, Non-dimensional cellular force scaled with possible parameters affecting the cellular force generation during confined migration for *D*_0_*/ϕ* = 1.67. **b**, Phase diagram depicting the zones of non-plastic and plastic nuclear deformation required for pore entry for different values of *E*_*n*_, *E*_*n*_*/E*_*T*_ and *D*_0_*/ϕ*. **c**, Proposed model of nuclear damage. Compressive forces imposed by the surrounding tissues cause initial nuclear membrane damage. This serves as the precursor to nuclear bleb formation.

## Discussion

The numerical model of cell migration under confinement presented in this study incorporates essential cellular features at the microscale, namely, nuclear elastoplasticity and viscoelasticity of other cellular components and extracellular matrices in addition to stress-stiffening of cytoplasm that make it more realistic than previous FE models (Supplementary Table 1). Our model predicts that while migrating through stiff matrices (*E*_*T*_ */E*_*n*_ > 1), the nucleus undergoes plastic deformation (Fig. 5b). Since Lamin A/C levels scale with tissue stiffness, our results of cells with stiff nuclei migrating through stiff tissues correspond to cancers such as osteosarcoma, wherein migration-induced DNA damage has been shown to cause genomic heterogeneity [35]. Stiff nuclei have been reported to result from increased lamin A concentration in the nucleus [36], especially in the genetic mutations caused in the Hutchinson Gilford Progeria Syndrome (HGPS) [37, 38]. In comparison, the absence of nuclear kinks in cells with soft nuclei passing through stiff matrices suggests that nuclear softening may represent a robust strategy utilized by cells to migrate through pores without facing nuclear membrane rupture. Consistent with this idea, *γ*H2Ax levels were comparable in control and Blebb-treated cells. The relative insensitivity of average cell speed to *E*_*n*_*/E*_*T*_ suggests that tissue stiffness-dependent temporal tuning of nuclear stiffness by lamin A/C phosphorylation may enable cancer cells to migrate at comparable efficiency through tissues of varying composition and pore sizes. Approximating the nuclear membrane and the lamina as a simply supported elastic plate of thickness *h*, the flexural rigidity (*F*_*D*_) can be given as [39]:

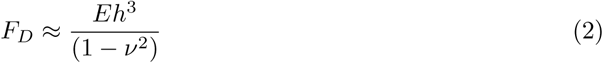

where, *E* and *ν* are elastic properties of the nuclear lamina. Since thickness *h* has a relatively greater influence on flexural rigidity compared to stiffness *E*, a thin lamina is more prone to damage due to bending than a soft lamina. This might explain increased blebbing due to nuclear damage reported in laminopathies (loss of lamin A/C) but not in immune cells or cancers with soft nuclei.

The chromatin contained within the nucleus is a major determinant of nuclear deformation. Mechanotransduction between the actin cytoskeleton and nucleoplasm through the interconnecting LINC complex has been shown to play a critical role in chromatin dynamics during DNA repair [40] and gene transcription [41, 42]. The spatial organization of the chromatin changes with nuclear stress and shape change. Compact chromatin network acts as an elastic spring to resist small deformations [42]. While small strains lead to strain stiffening of nuclei due to chromatin compaction [43], large deformation of nuclei is facilitated by the actin and vimentin cytoskeleton [44] and Lamin A/C [43]. A full rupture in the membrane allows the intranuclear pressure to become more than the intracellular pressure, thus facilitating the leakage of genetic material into the cytosol. An alternate mechanism is also observed in cells migrating under confinement where the nuclear membrane gets mechanically decoupled from the nuclear lamina which leads to membrane blebbing due to chromatin flow into this vacant pocket of space [45]. This experimental observation can be linked to our model prediction where we find that the nuclear lamina bends when the nucleus undergoes plastic deformation. This extreme nuclear bending as seen in Figs. 3b and 3d might result in delamination of the nuclear cortex from the nuclear membrane leading to bleb formation.

Nuclear blebbing has also been reported to be caused by influx of water into the nucleus under confinement [46]. In this study, we did not consider the effect of water influx into the nucleus, the characteristic time (*τ*) for which is found to be governed by the equation: 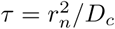, where *r*_*n*_ is the radius of nucleus and *D*_*c*_ is the diffusion coefficient of water. Thus, for an undeformed nucleus of *r*_*n*_ = 2.5 *µ*m and *D*_*c*_ = 50 *µ*m^2^*/*s [47], *τ* = 0.125 *s*, which is extremely small compared to the timescales of migration (minutes to hours), even smaller for deformed nuclei. Moreover, since we consider a quasi-static viscoelastic description of the model, we did not consider the transient poroelasticity of nuclei or cytoplasm. Our results show that there is a time-dependent spatial gradient of compressive forces on the nuclear lamina due to actomyosin fibres during nuclear entry into a pore. In such a situation, by the time the nucleus completely enters into the pore, the front of the nucleus relative to the direction of migration has been under compressive stresses much longer than the rear. Therefore, the probability of nuclear membrane rupture at the nuclear front is much higher than at the rear as has been consistently reported in experiments. Compressive stresses beyond a critical threshold (yield stress *σ*_*y*_) causes the nucleus to yield and this yield zone is also found to spread starting from the frontolateral region of the nucleus where necking occurs due to constriction to the rear of the nucleus (Supplementary Fig. 4b). In case of stiff nuclei (*E*_*T*_ ≈ 1 kPa) transiting through stiff matrices (*E*_*T*_ ≈ 2 kPa), plasticity-induced deformation of the nuclear membrane leads to buckling and finally membrane failure.

Nuclear membrane rupture in micropipette aspiration experiments has been attributed to tensile stresses at the anterior periphery of the nucleus [22, 48]. Similar results have been recapitulated by Cao et al. [15] in their model where they proposed that the front edge of the nucleus might be susceptible to tensile-stress induced damage during migration through ECM-like environments. Though these findings implicate tensile stresses as a factor contributing to nuclear damage, nuclei are primarily subjected to compressive forces during confined migration. Our observations of kink formation at the tip of the nuclear membrane proximal to the direction of migration correlates with experimental observations of the spatial location of nuclear damage during migration through extreme confinement in stiff environments [18, 19, 49]. The kink formation might be a consequence of excessive bending of the nuclear lamina driven by the combined effects of tissue stiffness and the peri-nuclear cytoskeleton. Formation of smaller lateral kinks might aid in the initiation of plastic deformation. We propose that rapid buildup of compressive and tensile stresses at the point of pore entry induces nuclear envelope damage; subsequent delamination of the lamina from the nuclear membrane may serve as a precursor to experimentally observed nuclear blebbing (Fig. 4c). The genetic material, already under significant external pressure and previously held back by the structural integrity of the nuclear lamina, then oozes out through the damaged orifice to form a bleb that may eventually rupture subject to membrane tension. However, unless the damage is extreme, nuclear rupture is repaired using ESCRT machinery [18, 20, 45]. Our results suggest that compressive stress-induced membrane damage and nuclear blebbing only occurs in a specific window depending on nuclear/tissue stiffness and extent of confinement, and may be critical for migration through stiff environments. Our experimental observations indeed support this idea as change in nuclear circularity indicative of plastic deformation of the nucleus was only observed in RO-treated cells, where DNA damage was maximum. The lack of plastic deformation in Bleb-treated cells which were more invasive and had lesser DNA damage suggests that nuclear softening may be a more effective invasion strategy compared to nuclear plasticity.

In conclusion, we have developed a numerical model of confined cell migration that contributes to our understanding of the underlying physics of nuclear deformation and stresses during confined migration. We further validate our key prediction of nuclear plasticity leading to nuclear damage using experiments wherein RO-induced nuclear stiffening led to plastic deformation and higher DNA damage. Our model suggests that nuclear membrane damage in stiff nuclei plastically deformed by compressive stresses, may serve as the precursor for bleb formation that ultimately facilitates successful migration of a cell through stiff tissues.

## Methods

### Computational Methods

For studying dynamics of confined cell migration, a plane strain finite element (FE) model of the system was created in ABAQUS^®^. FE models involve discretizing the system into smaller elements by meshing it (dividing the system into several discrete polygonal elements/parts). Numerical techniques (e.g., Runge-Kutta technique) are then used to arrive at an approximate solution to an equation of the general form [*K*]{*u*} = {*F*}. This is analogous to a Hookean spring with [*K*] being a matrix representing spring stiffness, {*u*} representing a displacement vector and {*F*} denoting a vector of applied force. This equation is computed at each node of each polygonal element that the object is made of (See Supplementary Information for further details). Discontinuities resulting due to cellular organelles and granular structures at the nanoscale are homogenized and considered as a continuum at the microscale.

A computational domain needs to be selected for numerical simulations in FEM so that boundary conditions (BCs) are applied to the PDEs that are solved as part of the problem. Since our desired direction of cell migration is the +*x*−direction, we assume that the deformation or volume change of the cell perpendicular to the plane of migration (*xy*−plane) would be much lower than that in plane and hence can be neglected. This is complemented by our chosen material parameters of all system components where the Poisson’s ratio (*ν*) is 0.3 indicating that all the materials are compressible (Supplementary Table 2), that is, a change in area in the *xy*−plane does not accompany a similar change in the *yz*− or *xz*−planes. This assumption reduces the complexity of the system from 3D to a 2D plane strain problem. This assumption also finds credibility in experimental observations of cell migration through microchannels where the cell deforms or gets polarized in the direction of migration but the accompanying lateral deformation is negligible [50, 51].

In our formulation, entry of a 10 *µ*m diameter cell with a 5 *µ*m diameter nucleus into a pore ({3, 4} *µ*m diameter) at the interface of two tissues was simulated with the system assumed to be in a quasi-static state for the entire duration of the simulation (Fig. S1a, b). This was verified by observing the internal energy of the system to be much lower than the kinetic energy. Pore entry was mediated by active protrusive forces generated by the cell at the cell front [52, 53]. A comparison of the salient features of two other FE models [15, 54] with our model is presented in Supplementary Table 1.

The cell is composed of the following components: cell membrane, cytoplasm, nuclear membrane and nucleus. All the components except the nucleus are approximated as Kelvin-Voigt viscoelastic elements [55] where stress developed in a system depends on the strain and strain rate and is given by the equation 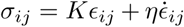. Here, *K* is an elasticity modulus that corresponds to spring or solid stiffness and *η* is viscosity of the constituent fluid. The subscript “*ij*” refers to the Einstein convention in the spatial dimension. The viscoelastic character of each component in the system is represented in the form of normalized creep compliance (Fig. S1c). Creep is a characteristic feature of a viscoelastic material that defines the amount by which a system deforms under persistent stress. Compliance is the reciprocal of stiffness (Pa) and is a measure of the ease with which a body deforms under stress.

The nucleus is considered to be elastoplastic with its behaviour described by a power law equation *σ* = *a* + *bϵ*^*n*^, similar to Ludwik’s equation [56]. The coefficients *a, b* and the exponent *n* were estimated to be equal to 41 Pa, 17 Pa and 2.89 respectively (*R*^2^ = 0.9981), with data trend similar to those reported by [28]. The elastic properties of the nucleus arise from the lamin network below the nuclear membrane and chromatin fibres. Plastic nature of nuclei have been reported in several studies demonstrating the irreversible change in shape of nuclei deformed under stress [28, 29, 43]. A recent study [43] also demonstrated that nuclei stiffen progressively with strain, a phenomenon attributed to chromatin compaction at small strains and to Lamin A/C at large strains. Plastic deformation in non-fibrous biological materials arise due to irreversible dislocation or dislodgement of molecules from their unperturbed positions. In fibrous biological materials like collagen, plasticity under tensile strains is caused due to unentanglement of fibers [12]. The tissues representing the two sides of the interface are also modelled to be viscoelastic. A Poisson’s ratio of *ν* = 0.3 which is a typical value considered for compressible biomaterials [15], was chosen for the cellular components as well as the two tissues. This value also sits well with our assumption that the out-of-plane volume change is negligible compared to the in-plane deformation. All elastic material properties are listed in Supplementary Table 2.

Cells are known to exert forces by actomyosin contraction resulting from myosin motors sliding on actin filaments [10, 11]. During migration, cells regulate F-actin polymerization and generate protrusions [57], leading to an increase in force generated. Crosslinking of actin filaments with proteins such as *α*-actinin, filamin and scruin stiffens these fibers and they bundle together to generate protrusive forces. Myosin II plays an important role in creation and regulation of stress fibers and force generation [57, 58]. To mimic these phenomena in our model, we assumed that the cell generates force (*F*_*P*_) at the leading edge in the direction of motion (x-axis) in a smooth monotonic fashion. *F*_*P*_ was assumed to be generated in a distributed fashion at the cell front as shown in Fig. S1d such that the magnitude of the maximum force (≈ 6.4 nN) generated by the cell remained in the physiologically relevant range [59]. In each step, after application of forces, if the cytoplasmic shear stress increased beyond 20 kPa, the cytoplasmic shear stiffness was increased in discrete steps from 1.0001 Pa to 1.1 Pa, and the system reequilibrated (Fig. S1e, Fig. S3) [30, 31, 60]. This variation can be curve-fit as per the equation *log*_10_(*E* − 1) = 49.41 *log*_10_*σ*_shear_ − 68.31, where *E* and *σ*_*shear*_ represent cytoplasmic stiffness and shear stress, respectively. The simulation was stopped once the nucleus enters the pore completely.

An explicit formulation was implemented to successfully resolve large nonlinear deformations in meshes in the Lagrangian or material domain. In this energy-based formulation, the stable time increment (Δ*t*) to solve the numerical problem depends on the stress wave velocity through the smallest element in the mesh (Δ*t* ≈ *L*_*min*_*/c*_*d*_), where *L*_*min*_ is the smallest element dimension in the mesh and *c*_*d*_ is the dilatational wave speed through the element. Mass scaling was used to ensure that Δ*t* was of the order O(−4). Frictionless hard contact was assumed at the cell-gel interface to simulate non-adherence of cell to the gel or channel walls. The cell surface was thus allowed to separate after contact with the gel surface. A total of 37436 bilinear plane strain CPE4R elements were used in the model, of which the two tissues were composed of 11895 and 11887 elements and the cell was composed of 13654 elements. The minimum element dimension was 0.01 *µ*m and the maximum was 20 *µ*m. The mesh size was modulated so as to be fine in the regions that were expected to come in contact or that would undergo large deformation. The cell membrane, 0.05 *µ*m in thickness, had 5 elements in the through-thickness direction to mitigate the effects of excess artificial bending stiffness of the membranes.

### Experimental Methods

#### Cell culture and reagents

HT-1080 fibrosarcoma cells obtained from National Center for Cell Science (NCCS) (Pune, India), were cultured in DMEM (high glucose, Invitrogen) containing 10% FBS (Hi-media). For nuclear stiffness experiments, cells were plated sparsely on glass coverslips coated with rat-tail collagen I (Cat # 3867, Sigma) at a coating density of 10 *µ*g*/cm*^2^. Cells were incubated with DMSO (i.e., vehicle), 1 *µ*M blebbistatin (Cat # B0560, Sigma) or 10 *µ*M RO-3306 (Cat # ab141491, Abcam) for 12 hours prior to probing with AFM.

#### Atomic Force Microscopy (AFM) and Imaging

For measuring nuclear stiffness, stiff tips (32 kHz, TR400PB, Asylum Research) with nominal stiffness of 120 pN/nm were used, with exact values of cantilever stiffness determined using thermal calibration method. Cells were indented towards the center right on top of the nucleus, and indentation data more than 2000 nm were fitted with Hertz model to obtain estimates of nuclear stiffness.

For transwell migration studies, 10^5^ cells were seeded on the upper chamber of 24 well plate cell culture inserts containing 3 *µ*m pores (Cat # 353096, Merck). The inserts were coated with rat-tail collagen I. For creating a gradient, the upper chambers were filled with plain DMEM supplemented with drugs and the lower chambers filled with DMEM containing 20% FBS. After 8, 18 and 28 hrs, cells were fixed with 4% PFA and then stained with DAPI for 45 minutes. After washing with PBS, membrane was cut and mounted on a glass slide using mounting media. Confocal z-stack images were acquired at 20x magnification using Scanning Probe Confocal Microscope (Zeiss, LSM 780) at identical exposure and gain settings. Images analysis and quantification was performed using Fiji-Image J software. Translocation efficiency was calculated using the equation 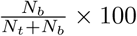 where *N*_*t*_ and *N*_*b*_ represent the number of DAPI-stained nuclei on the top and bottom surfaces of the membrane per frame.

For *γ*H2Ax staining, fixed cells were permeabilized with 0.1% Triton-X 100 for 8-10 mins, blocked with 2% bovine serum albumin (BSA) for 1 hr at room temperature, and incubated with *γ*H2Ax antibody rabbit monoclonal antibody (Cat # 9718S, CST) overnight at 4° C. The following day, after washing with PBS, Alexa-Fluor 555 anti-rabbit IgG was added for 2 hrs at room temperature. *γ*H2Ax images were acquired at 40x magnification using Scanning Probe Confocal Microscope (Zeiss, LSM 780) at identical exposure and gain settings.

## Acknowledgements

AM was supported by fellowship from IITB-Monash Research Academy. Authors acknowledge financial support from Department of Science and Technology (Govt. of India) (Grant # EMR/2016/005454). Authors would also like to thank IRCC, IIT Bombay for providing Bio-AFM and Confocal microscopy facilities.

## Author Contributions

AM and SS conceived the problem. AM, WY and SS were involved in formulating the problem. AM wrote the code and performed simulations with inputs from RS, WY and SS. AM, WY and SS were involved in the data analysis. AB performed the experiments with guidance from SS. AM and SS wrote the manuscript.

## Supplementary Information

### The Finite Element Method

The Finite Element Method (FEM) is a numerical technique that discretizes an object or structure with complex geometry into several small parts over which partial differential equations (PDEs) that pertain to the specific physics problem can be solved. A PDE (also referred to as strong form) describing the physical problem is converted to an integral of a lower differentiation order (weak form) which is then discretized into small elements. Thus, the integral or weak form gives way to a summation of the parts or elements to eventually result in a general set of simultaneous algebraic equation of the form:

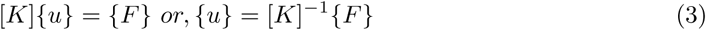

Eq. (3) is analogous to a Hookean spring with [K] being a matrix representing spring stiffness, {u} representing a displacement vector and {F} denoting a vector of applied force although it is general and can pertain to several physical systems. This equation is computed at each node of each polygonal element that the object is made of. All material constitutive models are converted into the form of Eq. (3) that guides the relation between stress and strain in the system.

### Viscoelasticity of cell and extracellular matrix (ECM)

Cell membrane (plasma membrane) and nuclear membrane are lipid bilayers that are dotted with various protein complexes and ion channels that allow for the transmigration of molecules. In a coarse-grained scenario, the cell membrane can be considered to be composed of a combination of a lipid bilayer, glycocalyx (polymer chains of glycolipids and glycoproteins [61]) and the actin cytoskeleton meshwork attached to the lipid bilayer. This composite cell membrane behaves as a viscoelastic material that flows like a viscous liquid over a short time duration but exhibits a solid-like elastic behaviour at sufficiently long timescales. This leads to the experimental observation of cells attaching themselves onto 2D substrates forming stable shapes [24–26]. A similar argument can be extended for the choice of the nuclear membrane and nuclear lamina composite as a viscoelastic material due to the similarities in their intrinsic composition with.

Considering the cell as a closed system consisting of a fibrous mixture (actin cytoskeleton, actomyosin fibres, microtubules and intermediate filaments) and a solvent (cytosol) with no net transport of molecules through the cell membrane, we model it as viscoelastic solid as opposed to poroelastic that assumes a net flux of solvent molecules. The tissue(s) through which the cell migrates is/are also considered as viscoelastic solids because we consider them to be individually closed systems which if stressed, lead to solvent molecules in the vicinity of the stressed region to get displaced from their initial locations temporarily before returning to their original position after stress is relieved. These assumptions are consistent with several experimental studies have demonstrated the viscoelastic nature of cells and tissues.

### Viscoelasticity formulation in the time-domain

To describe the constitutive relationship governing an isotropic viscoelastic material, we define the deviatoric and volumetric parts of the stress tensor. For the time-dependent deviatoric stress, time-varying shear strain *γ*_*dev*_(*t*) and shear stress *σ*_*dev*_(*t*) are related as:

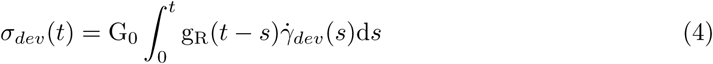

where *G*_0_ is the instantaneous shear modulus and *g*_*R*_(*t*) = *G*_*R*_(*t*)*/G*_0_ is the dimensionless time-dependent shear relaxation modulus of the viscoelastic material. The time-dependent volumetric behaviour (*σ*_*vol*_) of the material is defined as a change in hydrostatic pressure (*p*(*t*)) over time and is given by the equation:

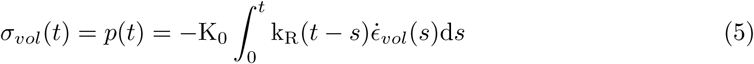

where *K*_0_ is the instantaneous bulk modulus and *k*_*R*_(*t*) = *K*_*R*_(*t*)*/K*_0_ is the dimensionless time-dependent bulk relaxation modulus of the viscoelastic material. The instantaneous moduli *G*_0_ and *K*_0_ are related to the Young’s modulus *E*_0_ and Poisson’s ratio *ν* as *G*_0_ = *E*_0_*/*2(1 + *ν*) and *K*_0_ = *E*_0_*/*3(1 − 2*ν*) respectively. A viscoelastic material is defined by a Prony series expansion of the dimensionless relaxation modulus given by the equation:

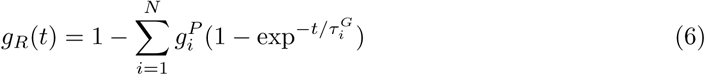

where N, 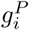 and 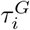, *i* = 1, 2,…, *N*, are material constants. The shear stress then is given by:

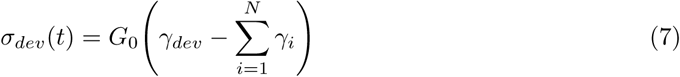

where 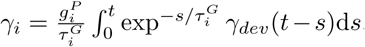. A similar expression can be acquired for the volumetric response, as shown below:

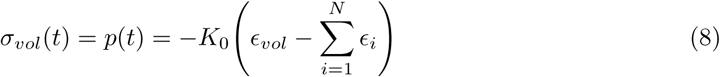

where, 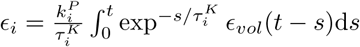.

### Plasticity of the nucleus

Previous studies have demonstrated that stressed nuclei undergo plastic deformation, i.e., they are irreversibly deformed under the application of stresses [28, 29]. Plastic deformation in non-fibrous biological materials arise due to irreversible dislocation or dislodgement of molecules from their unperturbed positions. In fibrous biological materials like collagen, plasticity under tensile strains is caused due to un-entanglement of fibers [12]. Plasticity is generally quantified as a strain or stress regime that extends beyond a critical threshold elastic limit below which molecular dislocations are reversible. An elastic material is assumed to have a linear stress-strain curve within a threshold termed as the proportional limit, beyond which the slope of the curve changes and the relation may become nonlinear. Plastic deformation leads to energy dissipation and therefore, the onset of plasticity signifies a new stable energy state for the material from the previous metastable strained state. Plasticity induced nuclear damage and rupture due to extreme stresses originating under confinement, for instance, may lead to genetic perturbation [18, 22].

### Cytoskeletal strain stiffening

Actin bundling proteins (ABPs) get attached to actin filaments with increasing stresses in the cytoplasm [30, 31]. Moreover, actin filaments frequently bundle together in a direction perpendicular to the direction of application of external force. These mechanisms contribute to the eventual stiffening of actomyosin networks. Studies indicate that depending on the actin concentration and crosslinking density the stiffness of such crosslinked fibres can change drastically [30–32]. In our model we implemented this experimental observation such that if the cytoplasmic shear stress increased beyond 20 kPa, the cytoplasmic shear stiffness was increased in discrete steps from 1.0001 Pa to 1.1 Pa, and the system re-equilibrated (Fig. S1e).

**Table 1:** Comparison with other FE-based cell migration models.

| Model Feature | Zhu and Mogilner [54] | Cao et. al. [15] | Current Study |
| --- | --- | --- | --- |
| Consideration of cytoskeleton | Y | N | Y |
| Cytoskeletal stiffening | N | N | Y |
| Nuclear Plasticity | N | Y | Y |
| Viscoelastic cellular components | N | N | Y |
| Viscoelastic gel/tissue/ECM | N | N | Y |

A comparison of the salient features of two other models [15, 54] besides our model is presented in Table S1, where ‘Y’ signifies the feature being accounted for in that model while ‘N’ implies absence of that feature. While this list is not exhaustive, it lists some of the mechanically and physically critical features that aid in cell migration in 3D matrices.

**Table 2:** Material parameters.

| Component | Density ($\rho$)<br>(kg/m<sup>3</sup>) | Young's<br>Modulus (E)<br>(Pa) | Poisson's<br>ratio ($\nu$) | Reference |
| --- | --- | --- | --- | --- |
| Cell Membrane | 1050 | 300 | 0.3 |  |
| Cytoplasm | 1030 | 1 | 0.3 | [62] |
| Nuclear Membrane | 1800 | 200-2000 | 0.3 | [63] |
| Nucleus | 1800 | 200-2000 | 0.3 | [43, 63] |
| Tissue 1, 2 | 1500 | 130-2000 | 0.3 | [64] |

## Supplementary Figure Legends

**Figure S1:**
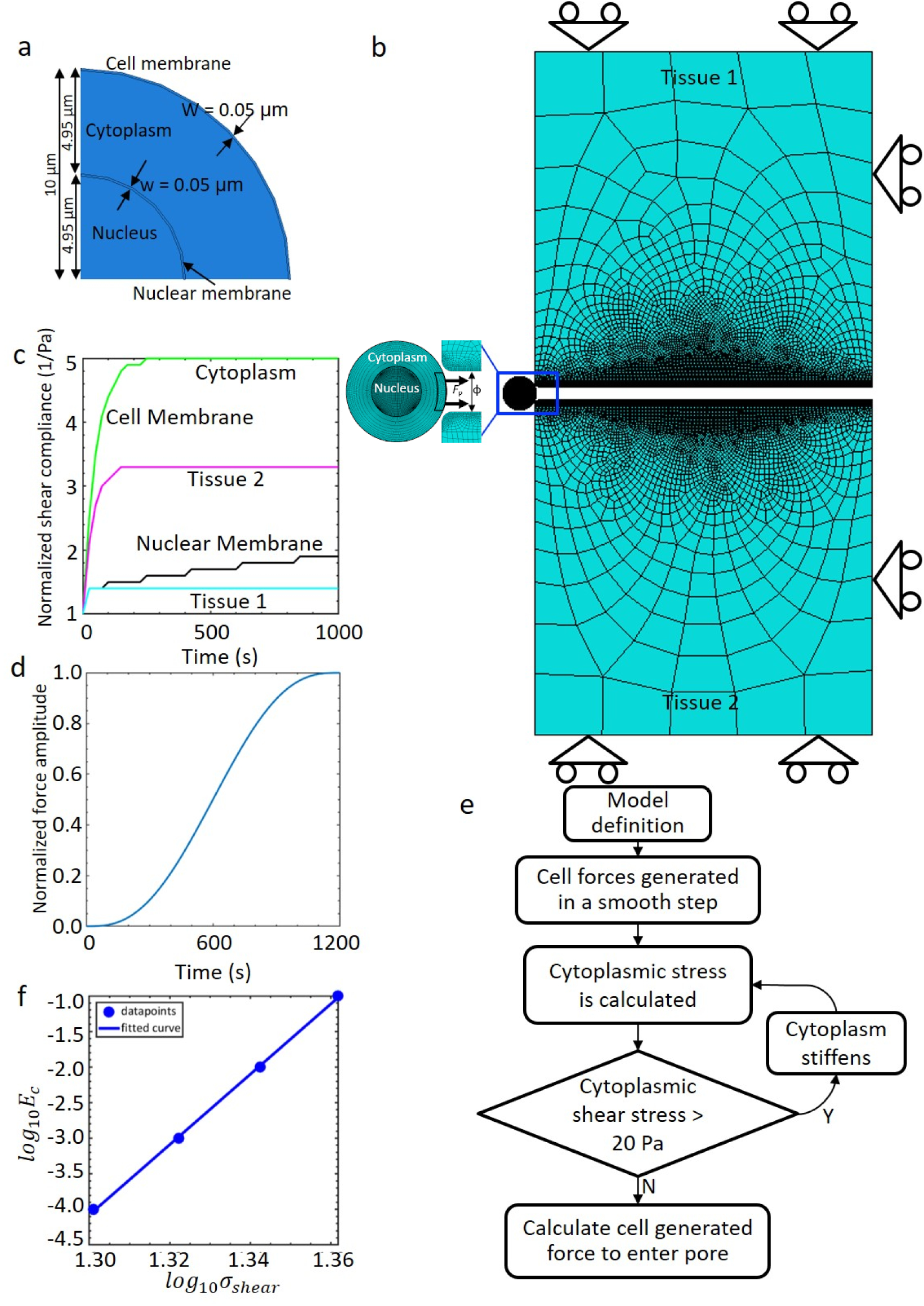
Model definition. **a**, Dimensions of various parts of the modelled cell. **b**, Finite element model with mesh. Lateral and transverse boundaries of the tissue (1 and 2) are constrained in their perpendicular directions. **c** Viscoelastic properties of various materials in the model. **d** Temporal variation of input force. **e** Simulation process flow. **f** Assumed dependence of cytoplasmic stiffness (*E*_*c*_) with shear stress (*σ*_shear_) encountered by the cell. *E*_*c*_ is increased in discrete steps as indicated by datapoints and a smooth curve is interpolated, i.e., the points are used to define a function between the two variables.

**Figure S2:**
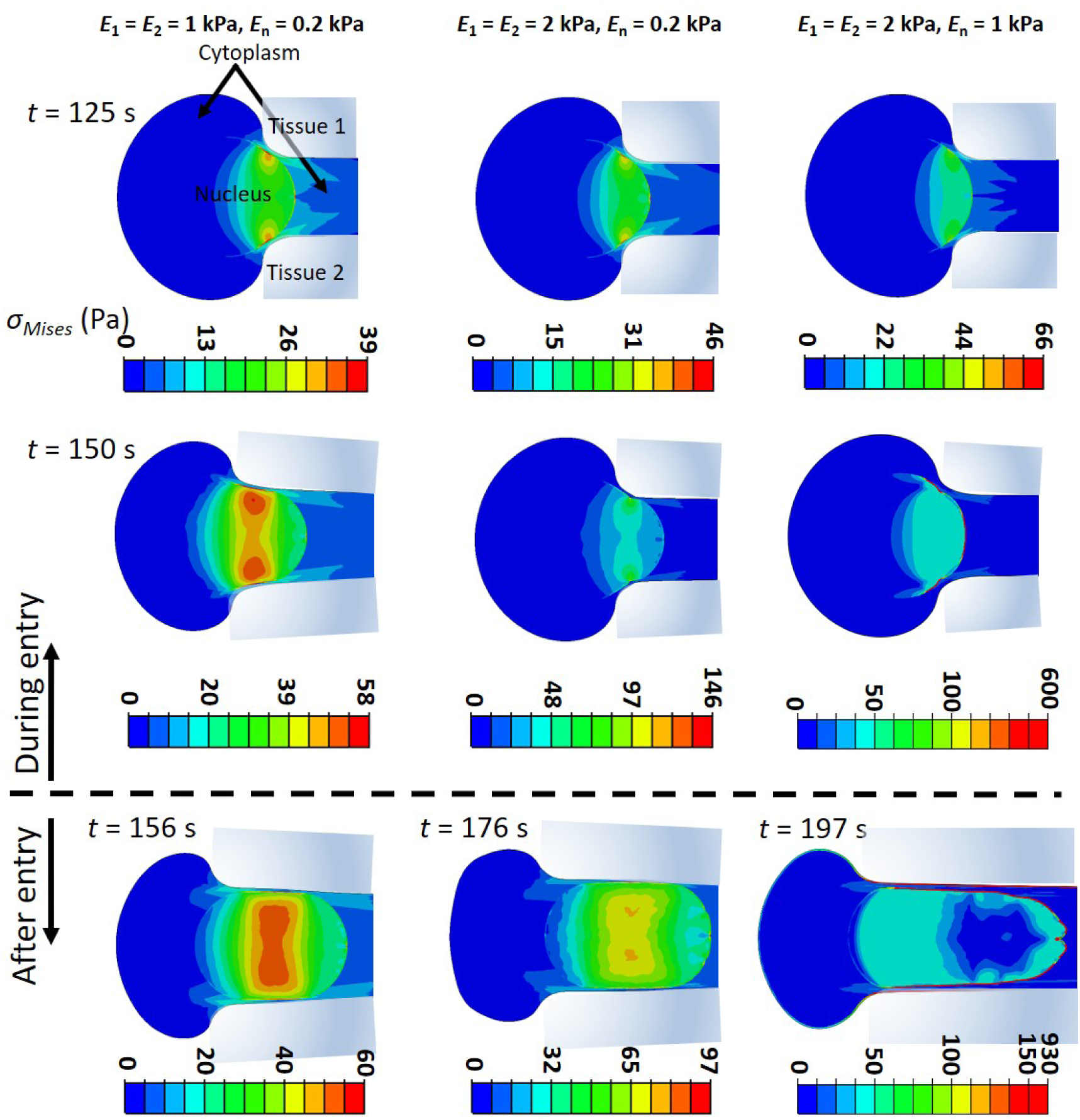
Stresses developed in the cytoplasm and the nucleus during pore entry. The spatiotemporal evolution of stress distribution during and just after entry of the 5*µ*m nucleus into a 3*µ*m pore, i.e., *D*_0_*/ϕ* = 1.67. Contours and colourbars indicate von Mises stresses developed in the cytoplasm and nucleus. The cell membrane has not been displayed in the figures for clarity.

**Figure S3:**
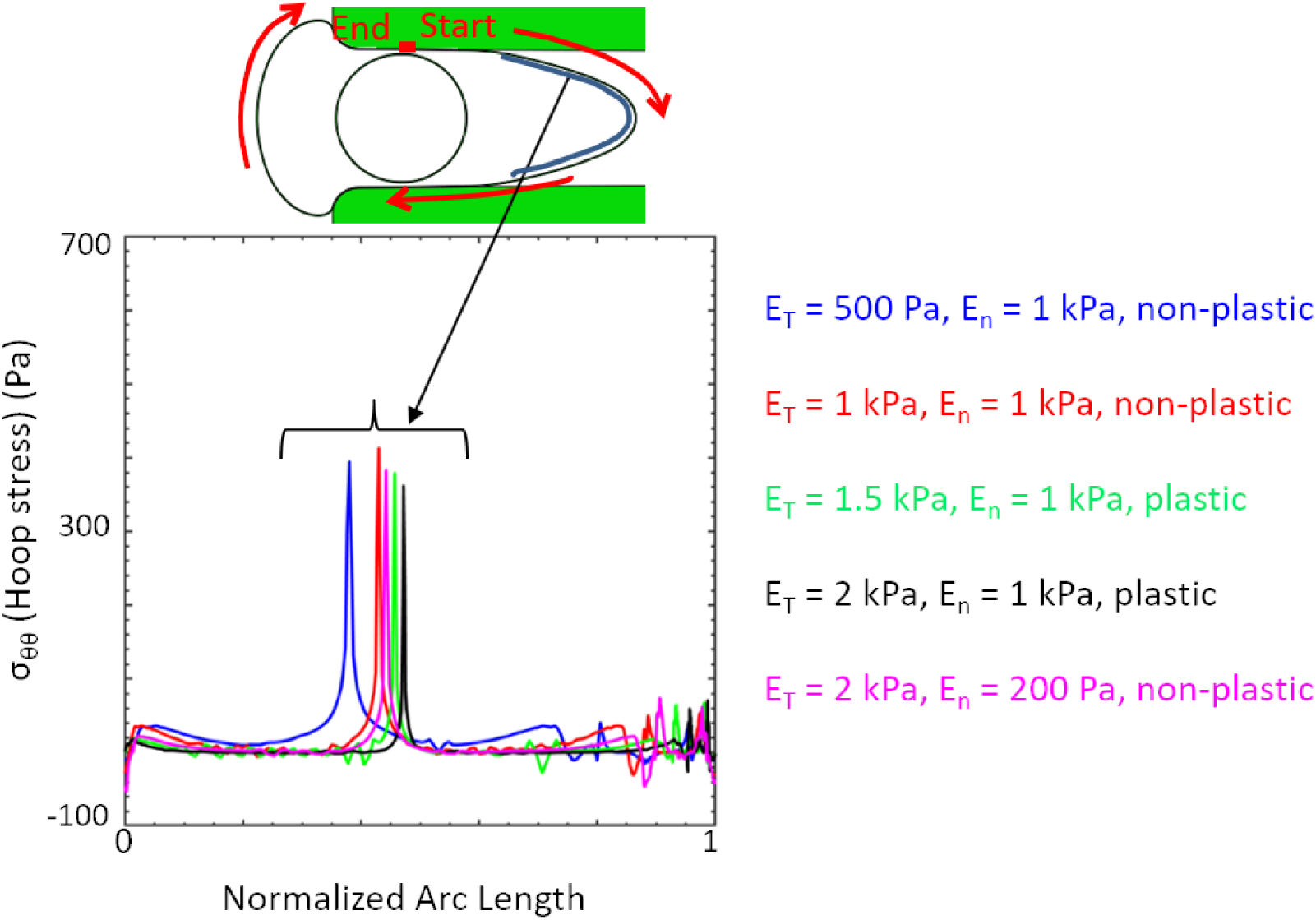
Hoop stresses developed in the cell membrane just after pore entry. For a confinement of *D*_0_*/ϕ* = 1.67, the spatial variation of hoop stresses along the length of the membrane for different combinations of *E*_*T*_ and *E*_*n*_.

**Figure S4:**
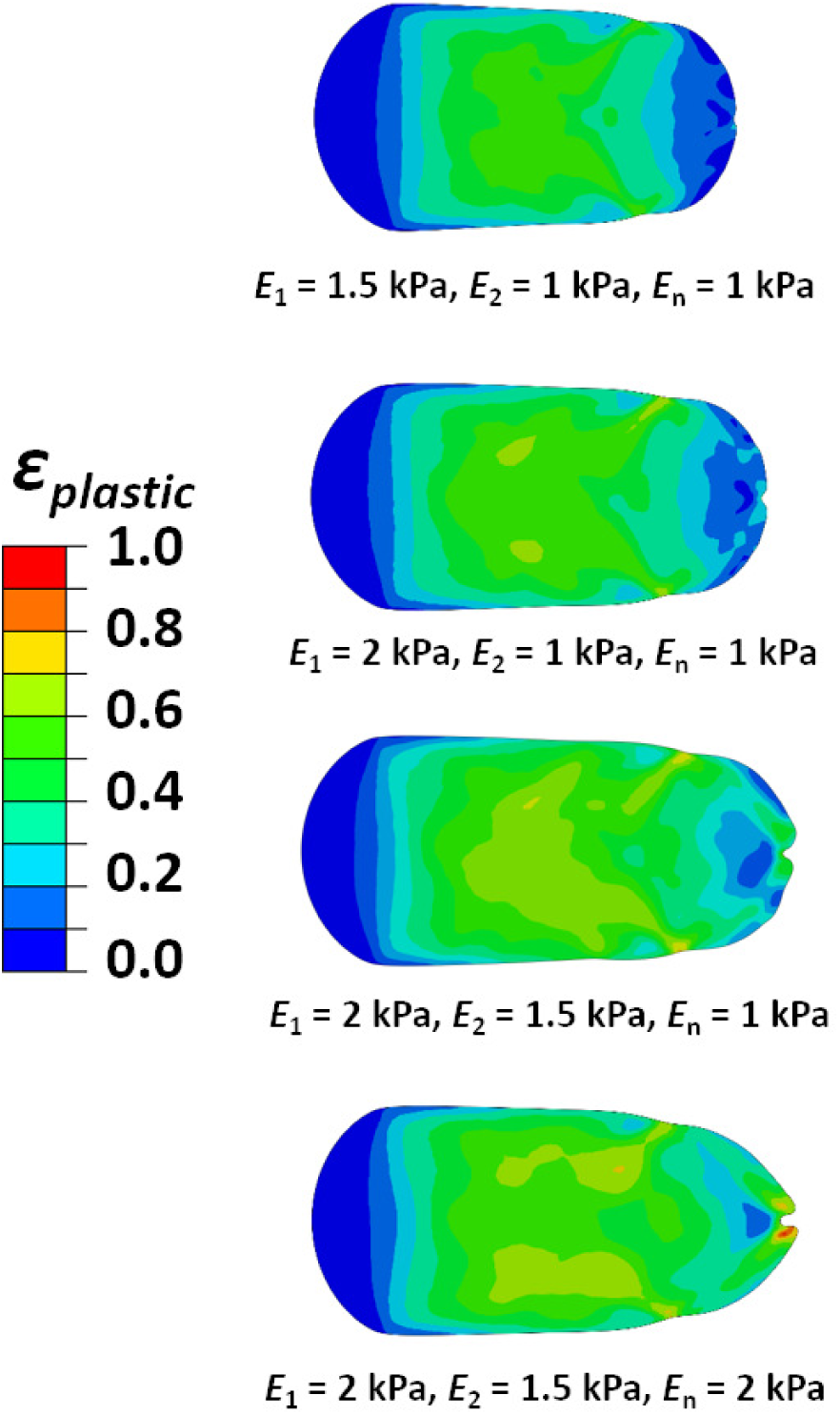
Plastically deformed nuclei in cells migrating through an interface. *E*_1_ and *E*_2_ refer to the Young’s moduli of tissues 1 and 2 on both sides of the interface. *D*_0_*/ϕ* = 1.67 for all the cases. Contours represent the spatial distribution of plastic strain.

**Figure S5:**
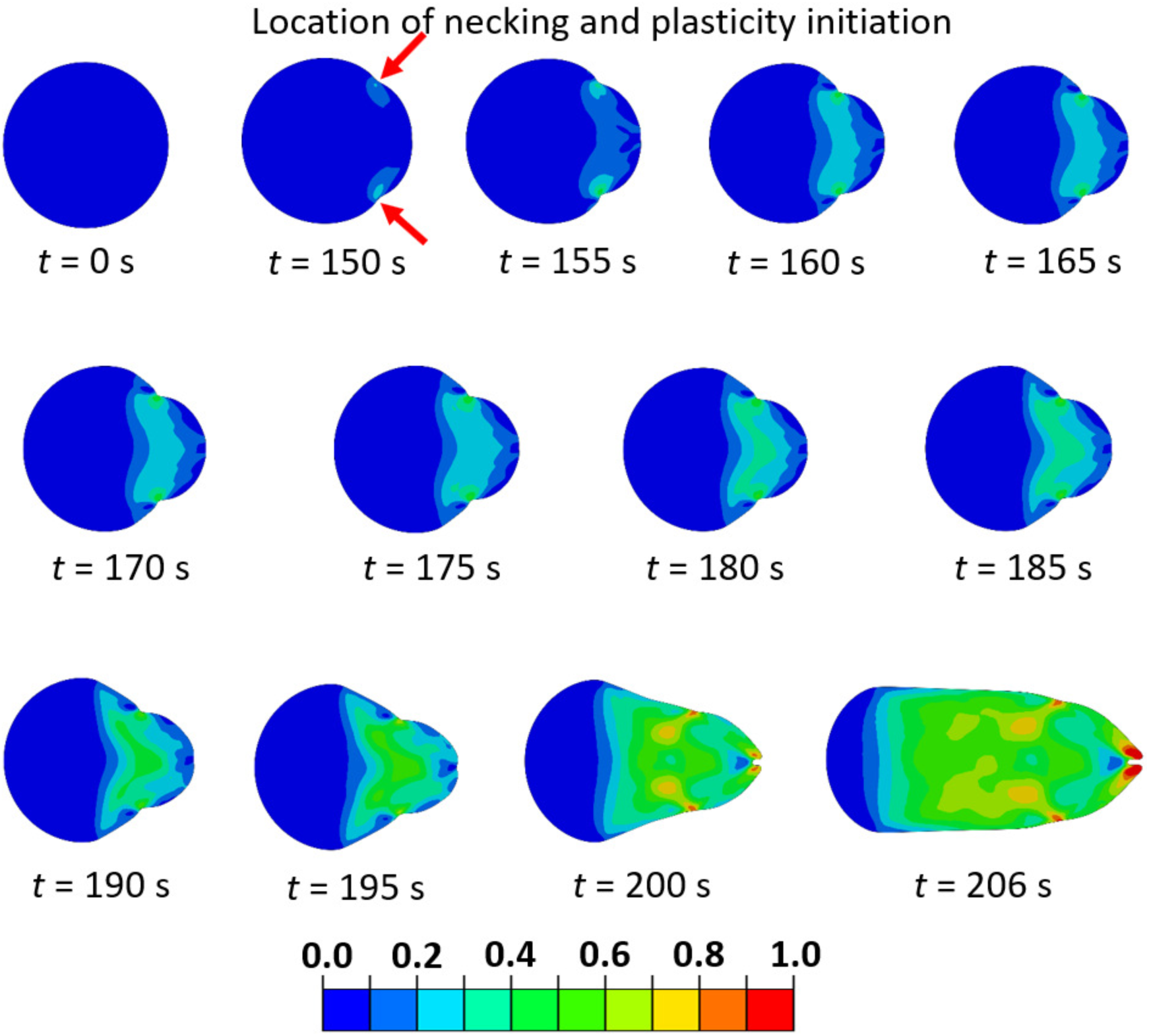
Spatiotemporal evolution of plastic deformation in stiff nucleus. Plastic strain accumulated in a cell as a function of time during constricted migration for *D*_0_*/ϕ* = 1.67. *E*_*n*_ = *E*_*T*_ = 2 kPa. Red arrows indicate the region where necking first occurs and plasticity is initiated. The colourbar indicates magnitude of plastic strain in the nucleus (*ϵ*_*plastic*_ = *ϵ*_*total*_ − *ϵ*_*elastic*_).

**Figure S6:**
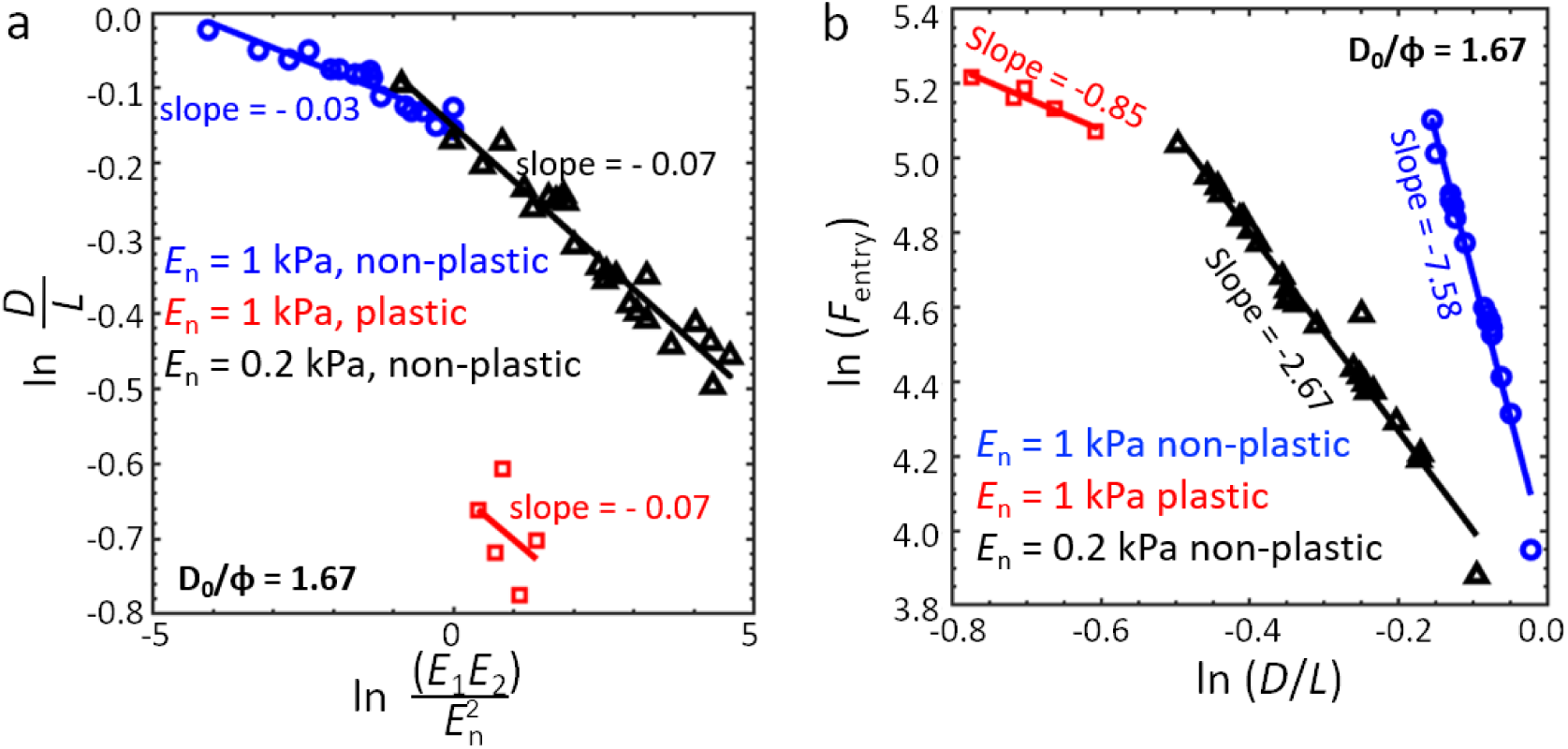
Scaling relationship of nuclear circularity. Scaling between nuclear circularity and **a** the coupled effect of tissue and nuclear stiffness, and **b** force required by a cell to enter a pore. All datapoints refer to the condition *D*_0_*/ϕ* = 1.67. *E*_1_ and *E*_2_ vary from 0.13 to 2 kPa.

